# A Quarter-Power Law for Translating Human Development into Equivalent Mouse and Rat Ages

**DOI:** 10.64898/2026.10.20.760430

**Authors:** Pablo Vázquez-Borsetti

## Abstract

Translating developmental time between laboratory rodents and humans is a central problem in preclinical research, yet no consensus mathematical framework exists. Linear models are known to be inadequate, and the logarithmic transformations conventionally used in the field do not fully linearize the relationship. Here we evaluated seven candidate functional forms relating human developmental age to rodent age, using a curated dataset of homologous developmental milestones in mouse, rat and human. Models were fitted by weighted nonlinear least squares and compared by leave-one-out cross-validation under a single model-independent weighting scheme derived from a LOESS variance estimate. Model specification was assessed using heteroscedasticity-robust RESET tests, which revealed significant residual curvature for the raw, logarithmic and log–log transformations, whereas the quarter-power transformation (*X* = Human^1*/*4^) was the only one leaving no detectable curvature in either species (mouse *F* = 0.90, *p* = 0.407; rat *F* = 1.83, *p* = 0.163). A quarter-power model with an additive constant (*y* = *bx*^1*/*4^ + *c*) achieved the lowest cross-validated error in mice and performed nearly as well as the best-performing model in rats, a free-power model with an additive constant, while using one fewer parameter. A third-power formulation, evaluated in light of the ongoing debate over metabolic scaling exponents, never ranked first in either species. Joint models incorporating a species dummy variable further supported a common quarter-power structure across both species, with species-specific coefficients. The additive constant was required in every formulation, indicating a systematic offset between developmental timescales that a purely scale-invariant power law does not capture. The resulting model provides a parsimonious, empirically supported framework for estimating equivalent developmental ages across humans, mice and rats.

## Introduction

Animal models are essential not only for advancing biological and medical knowledge, but also for translating fundamental discoveries into therapeutic interventions. However, successful translation critically depends on understanding both the similarities and differences between model species and humans. This challenge becomes even greater in developmental biology, where differences in the timing and progression of biological processes can lead to substantial divergence in phenotype and function across species. As a result, establishing developmental correspondence between model organisms and humans is a central issue in translational research. Among laboratory animals, rodents are the most widely used, making reliable frameworks to align their developmental trajectories with those of humans especially important.

Most comparative studies of animal development have focused on evolutionary patterns rather than on establishing quantitative frameworks for translating physiological and developmental processes between model organisms and humans. Although numerous studies have addressed this problem with varying levels of detail and rigor (B. Clancy et al. 2001; Barbara Clancy et al. 2007; Donaldson 1918; Dutta and Sengupta 2016; Finlay 2008; Hamdy et al. 2020; Romijn et al. 1991; Semple et al. 2013; Vargas Mamani 2020; Workman et al. 2013), no general consensus has emerged. There is still no widely accepted mathematical model to describe developmental relationships between humans, rats and mice, nor a standardized framework for interpreting such comparisons (Ortiz et al. 2022).

In previous work (Campos Eusebi et al. 2024), we showed that a linear model was insufficient to accurately describe these developmental relationships. Following the established tradition in this field, logarithmic transformations were applied to improve the fit. However, we also showed that this approach remained incomplete: although the transformed data were reasonably well fitted, they were not fully linearized, with a persistent residual convexity. This suggests that the underlying relationship may be inherently non-linear even after the transformations.

An alternative to logarithmic models that is ubiquitous in the description of natural phenomena is the power-law model, which describes the relationship between two variables as a power function. A well-known example in biology is Kleiber’s law, which states that basal metabolic rate scales with body mass to the power of 3*/*4. Consequently, larger organisms have a higher absolute metabolic rate but a lower metabolic rate per unit mass than smaller organisms. Power-law relationships of this kind are central to the study of allometry, which examines how biological traits, including body dimensions, physiological processes, and behavioral characteristics, scale with organismal size. In the context of biological growth, Huxley and Teissier formalized the use of power-law relationships to describe relative growth, with the exponent characterizing how two variables scale relative to each other (Huxley and Teissier 1936).

Power laws have also been used to describe patterns in the development of teeth and claws across vertebrates (Evans et al. 2021), early embryonic development in the fruit fly (Nikolić et al. 2024), and other biological processes (Stanley et al. 2000). Together, these examples illustrate how power-law relationships emerge across diverse biological processes and levels of organization, from metabolism and morphology to developmental dynamics. One key property of power-law models is scale invariance: the same relationship is maintained regardless of the scale at which the phenomenon is observed, so that changes in the magnitude of the variables do not alter the overall form of the relationship. When the independent variable is multiplied by a constant factor, the dependent variable changes by a corresponding factor determined by the scaling exponent. Moreover, power-law relationships become linear when both variables are expressed on logarithmic scales.

Evolutionary and developmental processes may constrain metabolic and geometric relationships in organisms, as developmental patterns appear to exhibit common features across biological scales and phenomena. This raises the question of whether developmental relationships could also be described by such laws. In this context, power-law models have been proposed to describe species’ lifespans and developmental timescales, which scale approximately with body mass to the one-quarter power (Lindstedt and Calder 1981).

In a companion study (Amici et al. 2026), we established the largest dataset to date of homologous developmental events across mouse, rat, and human. This comprehensive empirical framework enables precise cross-species comparisons and provides a robust reference for selecting animal models in developmental and translational research. However, the literature contains competing mathematical models and contradictory findings, making it difficult to establish which functional form best captures cross-species developmental relationships. Accordingly, the present study evaluates seven candidate functional forms relating developmental age across mouse, rat, and human. These include linear, logarithmic, and classical power-law models, as well as a modified power-law variant with an additive constant, all evaluated on this dataset.

## Methods

### Datasets

The methods used to obtain the datasets for this study are detailed extensively in a companion work on the comparative development of mice and humans (Amici et al. 2026) and a previous study on the comparative development of humans and rats (Campos Eusebi et al. 2024).

Briefly, literature searches were conducted to gather data on the comparative development of humans, rats, and mice. Data were retrieved using databases and search engines including NCBI/PubMed, Google Scholar, ResearchGate, and Elsevier/ScienceDirect, and organized into Excel spreadsheets, with Zotero used as the reference manager. Additional developmental milestones were incorporated, and formatting changes were implemented to ensure consistency with the dataset. The mouse dataset was derived primarily from Otis’s database (Otis and Brent 1954) and supplemented with an additional dataset generated by our team.

The compiled datasets included only information traceable to original empirical sources, excluding reviews or books, or any other source not traceable to primary research. Each database was curated and validated for accuracy and traceability. For instance, Otis’s database initially contained macaque milestones, which were excluded to focus on the comparative relationships between the two rodent species and humans.

The database structure included the following columns: *Species*; *Parameter*, indicating either a developmental milestone or a developmental process; age in days after fertilization (DAF); bibliographic source; observations; verbatim quotes; and DOI. The data were compiled and curated entirely manually, without the use of AI assistance.

### Data Processing and Analysis

The programs used for data processing, analysis, and visualization were developed in-house using Python. Scripts were written in Python 3.9 (Python Software Foundation) with the required libraries, including NumPy, Pandas, SciPy, Statsmodels, Scikit-learn, Matplotlib, and Seaborn. Software development was carried out primarily within the Visual Studio Code integrated development environment. Agentic assistance was used during code generation; however, all code was carefully reviewed, validated, and curated by the authors, who assume full responsibility for its accuracy and implementation.

### Candidate mathematical models

To describe the relationship between human DAF (*x*) and the response variable (*y*; rat or mouse DAF), we evaluated a series of candidate models representing different functional forms. These included logarithmic and power-law models with either fixed or freely estimated exponents, with and without an additive constant. Model parameters were estimated using weighted nonlinear least squares (WNLS). Observation weights were defined as inversely proportional to the estimated variance to account for heteroscedasticity in the data. Parameter estimation was performed using the Levenberg–Marquardt algorithm. Model residuals and influence measures were examined to assess model fit and the robustness of the parameter estimates. The candidate models were defined by the following equations:

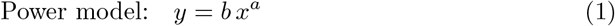

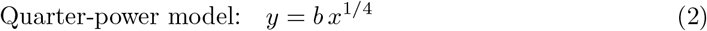

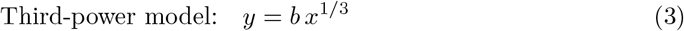

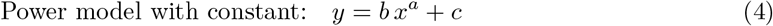

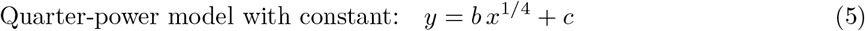

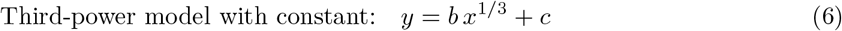

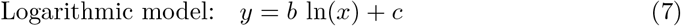

### Models incorporating dummy variables

To identify the mathematical model that best describes the developmental trajectories of both species simultaneously, each candidate model was extended by incorporating a binary dummy variable (*D*), where *D* = 0 for mice and *D* = 1 for rats. This approach enabled both species to be fitted jointly while allowing species-specific values for the scaling coefficient, additive constant, and, when appropriate, the exponent. Comparison of these joint models allowed us to determine whether a common mathematical relationship could adequately describe the developmental trajectories of both species and to identify the best-fitting formulation. The joint models were therefore defined as follows, with corresponding formulations without the additive constant also evaluated (Equations 13–16):

Logarithmic model:

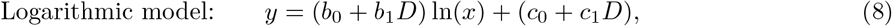

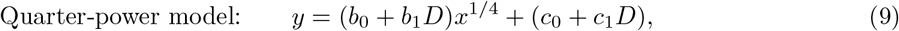

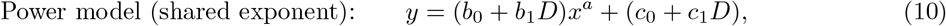

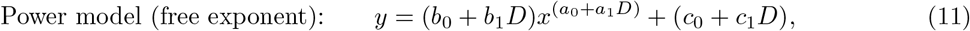

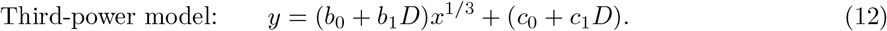

### Model comparison

To compare the models against each other, neutral weights were used for the WLS, which were generated based on the data smoothed using LOESS. Additionally, the LOESS was truncated to avoid edge effects. Predictive performance was assessed using leave-one-out cross-validation (LOOCV). For each candidate model, one observation was excluded, the model was fitted to the remaining observations, and the excluded observation was predicted. This procedure was repeated until every observation had served once as the validation sample. Model performance was quantified by the weighted root mean square error (WRMSE) of the cross-validated predictions, and the model with the lowest LOOCV-WRMSE was selected as the best-fitting model. The presence of linearity in the models was assessed through residual analysis and diagnostic tests robust to heteroscedasticity, specifically the RESET test with an HC3 covariance matrix. This approach is necessary because conventional specification tests assume homoscedasticity; when heteroscedasticity is present, as previously detected using the White test, conventional standard errors may lead to incorrect inference and inflated Type I error rates. Therefore, the robust RESET test was used to assess model specification and determine whether a nonlinear structure persisted in the residuals, thereby providing a criterion for identifying which transformations contributed to linearizing the observed relationships.

### Data Accessibility

The complete analysis code is available on GitHub at https://github.com/Vazquez-Borsetti/quarter-power-C-human-rodent-translation. The datasets, provided in CSV format, are accessible through both Zenodo (Pablo 2026) and the companion study’s GitHub repository at https://github.com/Vazquez-Borsetti/mouse-rat-human-development, and adhere to the FAIR (Findable, Accessible, Interoperable, and Reusable) principles. A more detailed description of the mouse dataset is available in the companion work (Amici et al. 2026). A subset of these data was previously published in (Campos Eusebi et al. 2024) and is available on GitHub at https://github.com/Vazquez-Borsetti/rat-and-human-comparative-development.

## Results

### Developmental relationships between rodents and humans

Mouse–human and rat–human developmental comparisons revealed remarkably similar patterns, with both species exhibiting a divergent and clearly non-linear relationship with human developmental age (Figure 1a–c). To further characterize the functional structure of these relationships, several transformations commonly used in the comparative developmental literature were evaluated to determine whether they reduced violations of the assumptions of homoscedasticity and linearity and enabled the establishment of linear correspondences. Visual inspection indicated that neither the single-log nor the log–log transformation fully linearized the relationships. The single-log transformation (Figure 1d–f) reduced the apparent curvature, whereas the log–log transformation (Figure 1g–i) provided a further improvement but still failed to achieve linearity, particularly during the early developmental period, where most homologous events are concentrated. Taken together, these findings suggest that the developmental relationships between rodents and humans are not fully captured by simple linear relationships, either on the original scale or after conventional logarithmic transformations. Although logarithmic transformation improved the linearity of the relationships, some curvature remained, indicating that alternative transformations or functional forms may provide a better representation of the developmental relationships. These results motivated the evaluation of alternative nonlinear models to identify a mathematical formulation capable of accurately describing the developmental correspondence between species.

**Figure 1:**
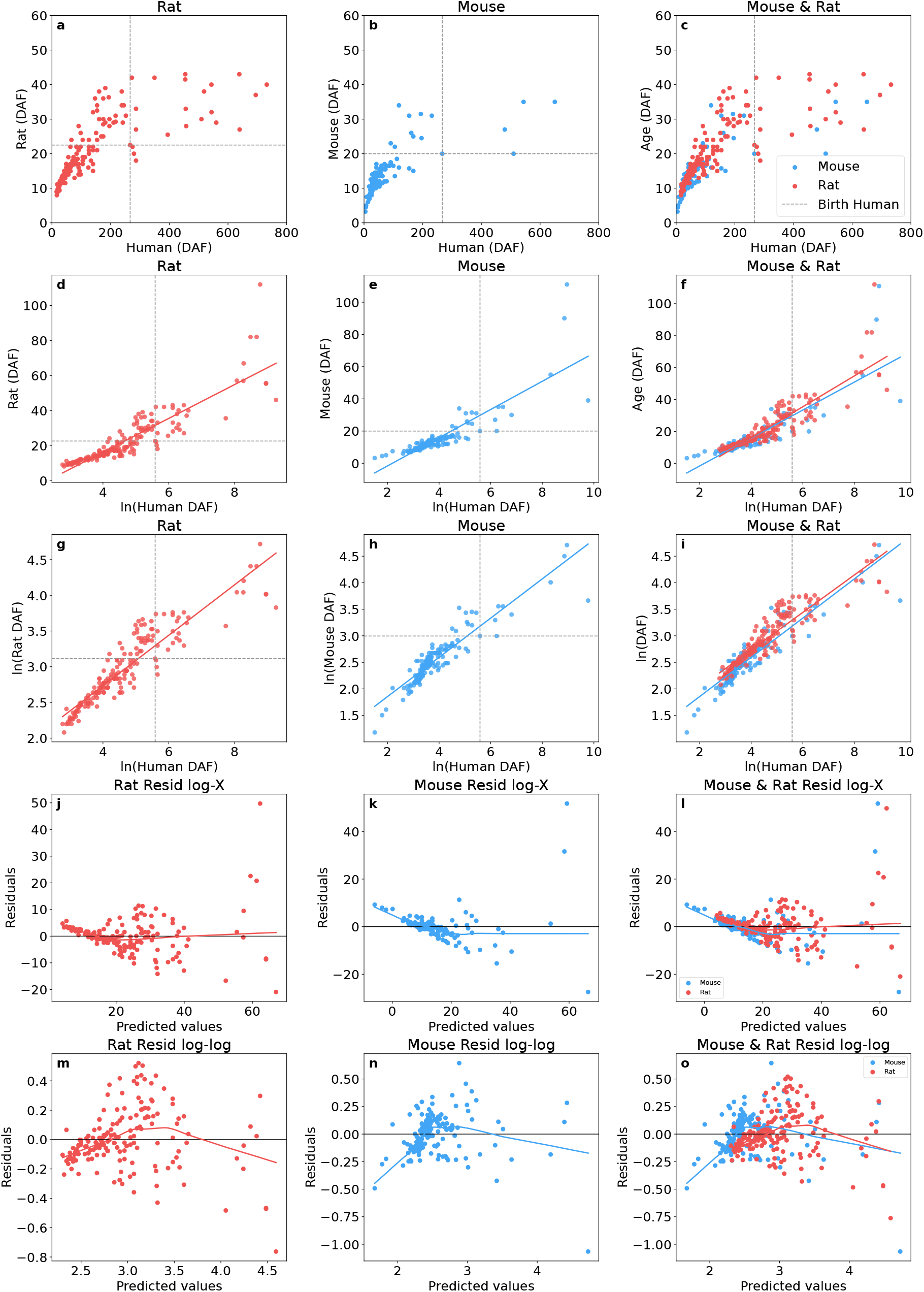
Developmental relationships between rodents and humans under different transformations. (a) Mouse–human relationship in the original scale. (b) Rat–human relationship in the original scale. (c) Overlay of the mouse–human and rat–human relationships in the original scale. (d–f) Corresponding single-log transformations. (g–i) Corresponding log–log transformations used to assess whether linearization improved the fit. (j–o) Residual plots for the different transformations. Although the transformations reduced the apparent curvature, substantial departures from linearity remained.

### Quarter-power model with an additive constant provides the best predictive performance

Seven nonlinear models were evaluated independently for rats and mice using WNLS and LOOCV (Table 1). Overall, models that included an additive constant outperformed their corresponding formulations without a constant. In both species, the quarter-power model with an additive constant demonstrated excellent performance. In mice, it achieved the lowest CV-WRMSE, making it the best-performing model for out-of-sample prediction. In rats, it ranked second, although the difference from the power model with a freely estimated exponent was only 0.23% (3.1826 vs. 3.1752), trailing behind a less parsimonious model such as the standard power model. Conversely, power models lacking an additive constant and the logarithmic model consistently yielded higher CV-WRMSE values, making the quarter-power model the only one to show consistent performance across both species.

**Table 1:**
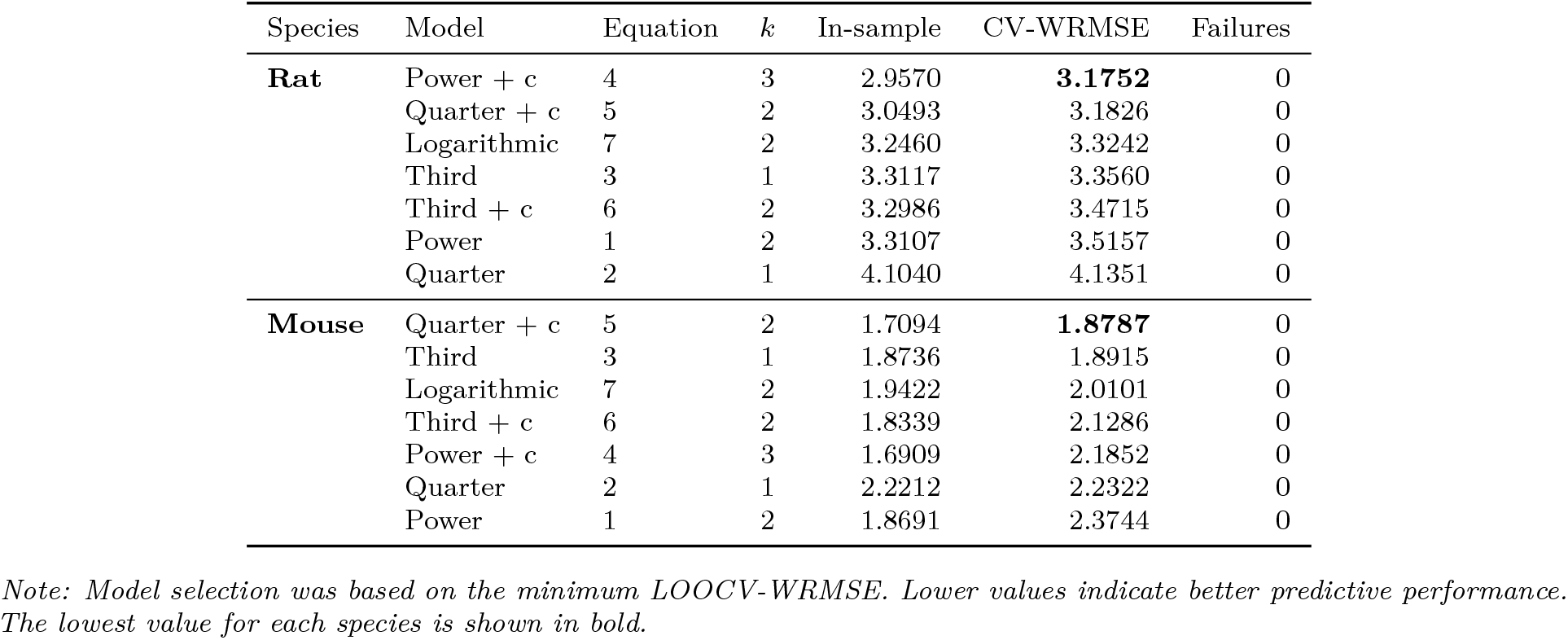
Leave-one-out cross-validation comparison of models within each species.

The difference between the in-sample WRMSE and the out-of-sample CV-WRMSE was relatively small for all models, suggesting limited overfitting and good generalization to unseen observations.

As shown in Figure 2, the quarter-power model with an additive constant (green) provided a fit that was visually very similar to that of the unconstrained power model with an additive constant (black). This similarity was particularly evident for rats, for which the two curves nearly overlapped across the observed range. The logarithmic model (purple), in contrast, showed a poorer fit, particularly toward the extremes of the observed range, where deviations from the data were more pronounced.

**Figure 2:**
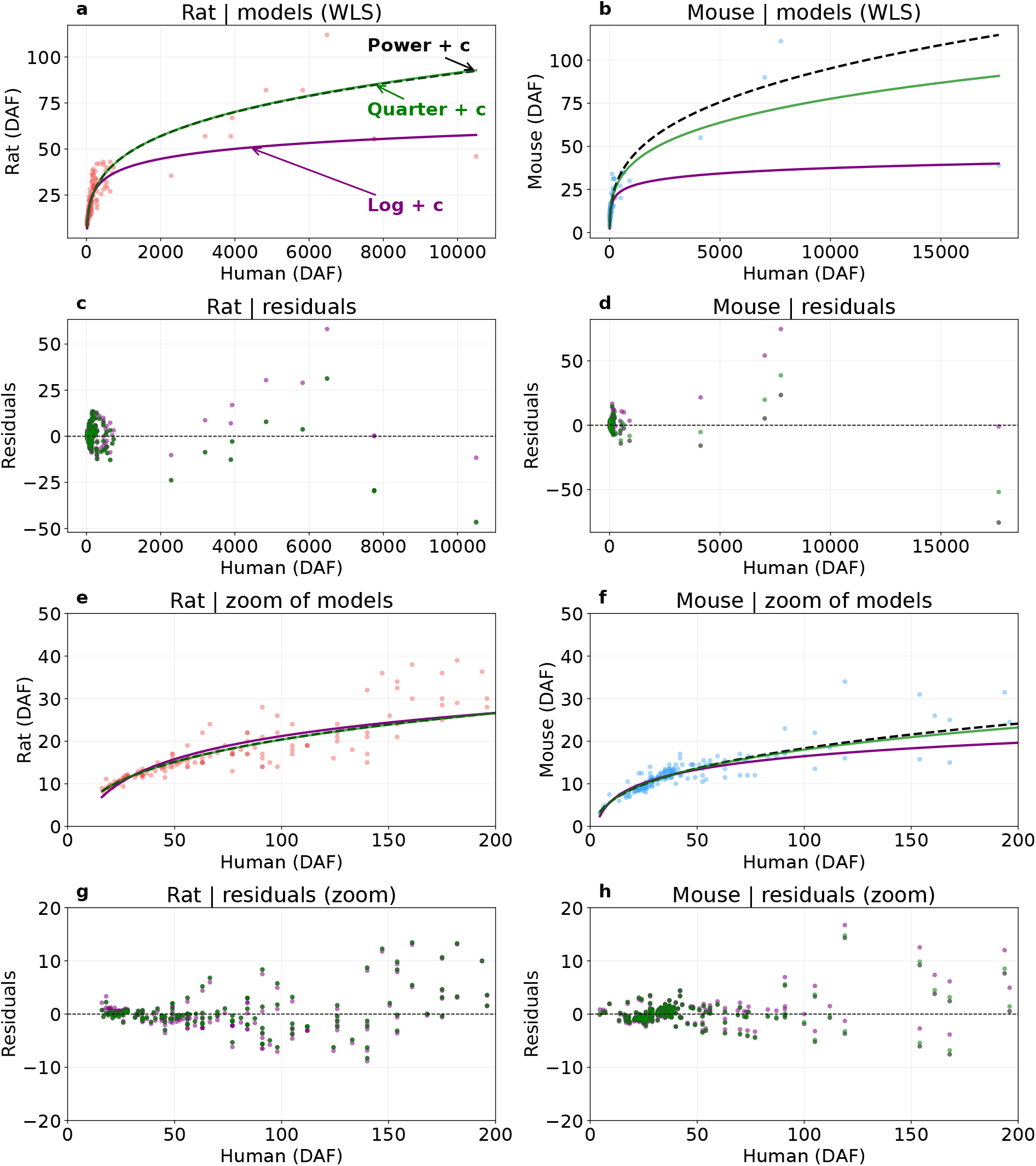
Comparison of weighted nonlinear models relating human and rodent developmental age, expressed as days after fertilization (DAF). WNLS fits for rats (a) and mice (b) comparing the power model with constant (*y* = *bx*^*a*^ + *c*; black), quarter-power model with constant (*y* = *bx*^1*/*4^ + *c*; green), and logarithmic model (*y* = *b* ln(*x*) + *c*; purple). Panels (c,d) show the residuals of the fitted models across the full developmental range. Panels (e,f) provide an enlarged view of the early developmental period, where the majority of homologous developmental events are concentrated, whereas panels (g,h) show the corresponding residuals for the same interval.

Table 2 presents the parameter estimates for each model fitted separately to the two species.

**Table 2:**
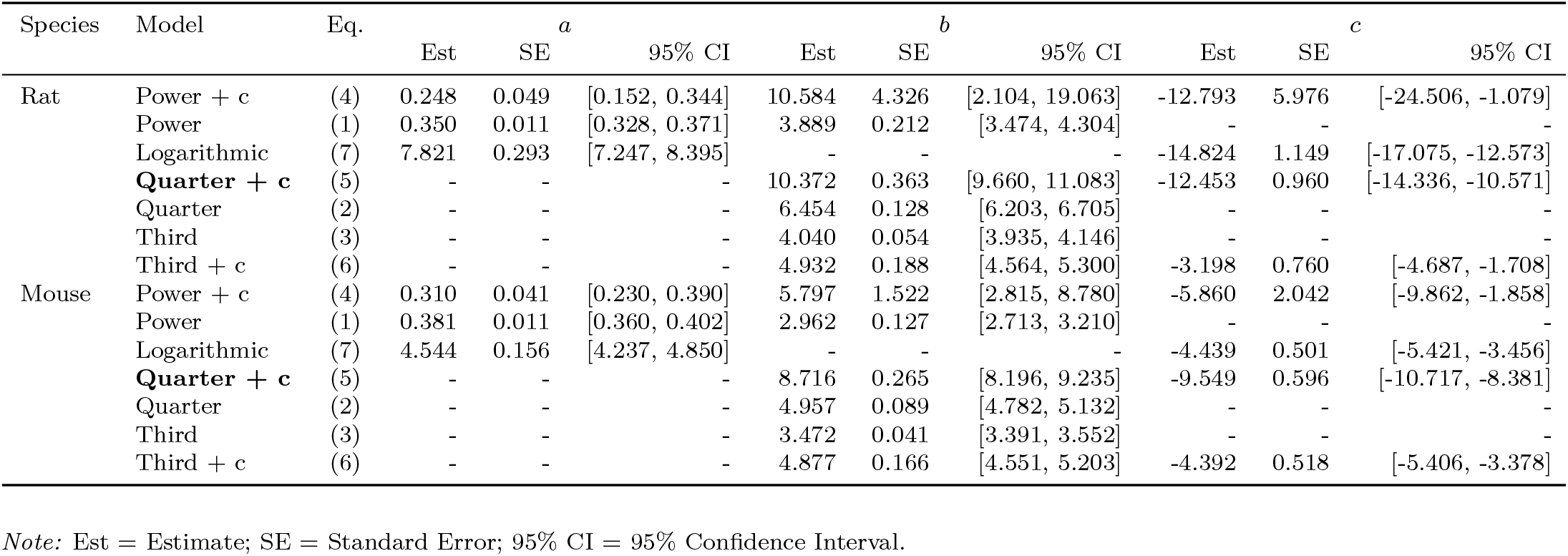
Estimated model parameters for each species.

Taking into account the arguments presented previously, along with the evidence that will be presented later, it was decided that the one-quarter power-plus-constant model was the best model to explain the developmental relationships between rodents and humans. Therefore, species-specific parameter estimates were selected for translating developmental age in rats and mice to the corresponding human developmental age. These parameter estimates are therefore also used in the companion study (Amici et al. 2026).

The transformations of human developmental age that allowed the human–rodent developmental relationships to be adequately linearized were assessed. The available transformations for linearizing the data (Raw, Log, Log–Log, and Quarter) were evaluated using WLS regression. Linearity was assessed using the RESET test with an HC3 heteroscedasticity-robust covariance estimator, given the presence of heteroscedasticity in the datasets. The robust RESET test showed significant evidence of residual curvature for the Raw transformation (mouse: *F* = 7.50, *p* = 0.0007; rat: *F* = 13.79, *p <* 0.0001), as well as for the Log transformation (mouse: *F* = 10.04, *p* = 0.0001; rat: *F* = 14.94, *p <* 0.0001) and the Log–Log transformation (mouse: *F* = 9.07, *p* = 0.0002; rat: *F* = 13.58, *p <* 0.0001). Thus, none of these three transformations fully linearized the allometric relationship between species. In contrast, the Quarter transformation (*X* = Human^0.25^) was the only transformation for which no detectable residual curvature remained: the robust RESET test was non-significant for both mouse (*F* = 0.90, *p* = 0.407) and rat (*F* = 1.83, *p* = 0.163). The White test confirmed significant residual heteroscedasticity across all transformations (*p <* 0.0001). Overall, these results indicate that the quarter-power transformation provided the most effective linearization of the human–rodent DAF relationships under the WLS framework.

### Combined models with species dummy variables

Combined models incorporating species dummy variables showed that developmental trajectories in mice and rats could be described by a common quarter-power relationship with human development. The quarter-power model including a constant term provided the best overall performance, while species-specific parameters accounted for differences between the two species (Table 3).

**Table 3:** Leave-One-Out Cross-Validation Comparison of Joint Models Including Species as a Dummy Variable.

| Model | Eq. no. | $k$ | In-sample | CV-WRMSE | Failures |
| --- | --- | --- | --- | --- | --- |
| Power (quarter) + c | 9 | 4 | 1.7826 | **2.9706** | 0 |
| Power (free) + c | 11 | 6 | 1.6949 | 3.1207 | 0 |
| Logarithmic | 8 | 4 | 2.0459 | 3.1777 | 0 |
| Power (shared) + c | 10 | 5 | 1.5885 | 3.2195 | 0 |
| Third-power + c | 12 | 4 | 2.1250 | 3.2347 | 0 |
| Third-power no constant | 13 | 2 | 2.1959 | 3.2390 | 0 |
| Power (shared) | 15 | 3 | 2.5579 | 3.3965 | 0 |
| Power (free) | 16 | 4 | 2.5384 | 3.4447 | 0 |
| Power (quarter) | 14 | 2 | 3.9664 | 3.8995 | 0 |

Note: Model selection was based on the minimum LOOCV-WRMSE. Lower values indicate better predictive performance. The lowest value for each species is shown in bold.

In most models, the coefficients associated with the species dummy variables were statistically significant, indicating that mice and rats differed in one or more model parameters. Although such differences are expected given the distinct developmental trajectories of the two species, incorporating species-specific parameters improved model fit without altering the underlying mathematical structure. This approach therefore allowed us to assess whether a common functional formulation could simultaneously describe developmental trajectories in both species while accounting for species-specific differences (Table 4).

**Table 4:** Estimated coefficients of joint models including species dummy variables.

| Eq. | Name | $b_0$ | $b_1$ | $c_0$ | $c_1$ | $a_0$ | $a_1$ |
| --- | --- | --- | --- | --- | --- | --- | --- |
| 8 | Logarithmic | 4.7744 *** | 2.7911 *** | -5.1676 *** | -8.5184 *** |  |  |
| 9 | Power (quarter) + c | 8.726 *** | 2.1118 *** | -9.5609 *** | -3.9878 *** |  |  |
| 10 | Power (shared) + c | 5.7095 *** | 1.0041 * | -5.6741 *** | -1.9269 ns | 0.3115 *** |  |
| 11 | Power (free) + c | 5.7913 *** | 2.859 ns | -5.7909 ** | -4.6869 ns | 0.3095 *** | -0.0314 ns |
| 12 | Third-power + c | 4.8655 *** | 0.6596 ** | -4.3485 *** | -0.9813 ns |  |  |
| 13 | Third-power no constant | 3.4582 *** | 0.6154 *** |  |  |  |  |
| 14 | Power (quarter) | 4.9292 *** | 1.3783 *** |  |  |  |  |
| 15 | Power (shared) | 2.9305 *** | 0.4226 *** |  |  | 0.3841 *** |  |
| 16 | Power (free) | 2.8364 *** | 0.5606 ** |  |  | 0.3928 *** | -0.0115 ns |

Note: Significance levels: \* $p < 0.05$ , \*\* $p < 0.01$ , \*\*\* $p < 0.001$ , ns = not significant. Species dummy variable: Rat = 1, Mouse = 0.

In a small number of cases, the coefficients associated with the dummy variables were not statistically significant, particularly the coefficient for the exponent (*a*_1_) in the free-exponent power model, both with and without a constant term. This finding is consistent with the hypothesis that interspecific differences can be captured by other species-specific parameters while maintaining a common scaling exponent. These results are also consistent with the use of a quarter-power exponent as a shared parameter across species. Moreover, the species-specific constant term (*c*_0_) was statistically significant in all models, indicating that its inclusion consistently improved model fit and predictive performance.

## Discussion

In a previous study on the comparative development of humans and rats (Campos Eusebi et al. 2024), it was shown that a linear relationship did not adequately describe the data, while logarithmic transformations, although providing an improvement, could not fully capture their convexity. In mice, this limitation was demonstrated in the first part of the present study. Found throughout nature in general, and in allometry in particular, power-law models are widely used. A key property of these models is that they can be linearized by applying logarithmic transformations to both variables. This mathematical property is one of the reasons why such transformations have been routinely used in allometric analyses. In this type of relationship, one variable scales as a power of the other. As a result, a relative change in one quantity produces a proportional relative change in the other, driven by a constant exponent. This scaling relationship remains consistent across different magnitudes of the variables, reflecting what is commonly referred to as scale invariance. These relationships have proven powerful for modeling physiological processes across species and have even been proposed to account for variation within species (Van Valkengoed et al. 2025). One classic example is the allometric relationship between body size, metabolic rate, and biological time. According to Kleiber’s law (Kleiber 1947), metabolic rates scale approximately to the 3/4 power of body mass, meaning that small animals exhibit higher oxygen consumption per unit of mass than larger animals. Such scaling relationships suggest that different biological processes might be organized by non-linear power laws that are conserved across different scales. Whether developmental timing follows comparable scaling relationships across species of different body sizes is an open empirical question. The present study addressed this question by fitting power-law models to cross-species age-translation data. Initial power-law models without an additive constant (Equation 1), however, failed to outperform logarithmic models. To improve the fit, a constant term was subsequently introduced. The freely estimated exponent was 0.248 for rats and 0.310 for mice. Despite the inclusion of the constant term, the exponent remained consistent with a quarter-power scaling relationship, with 1/4 falling within the confidence intervals for both species. Such scaling patterns are a recurrent feature in allometry, blood circulation times, embryonic growth and development, and lifespan increase proportionally to the fourth root of body mass (West et al. 1997). This same model, without the constant term, was evaluated (Equation 2). As previously observed in the free-power model, models lacking a constant term perform poorly, particularly when fitting the earliest developmental data. It is evident that there is an offset, at least when evaluating models across the entire developmental period, including late-stage data. The necessity of this negative additive constant (c) carries important biological implications. A pure quarter-power law (*c* = 0) assumes strict geometric scale invariance across the entire developmental timeline. The persistent residual convexity observed in the log-log transformations, together with the superior fit of Equations 4 and 5, may therefore indicate a developmental feature not captured by the conventional power-law formulation. Although pure power laws are widely used to describe allometric relationships, their biological basis and applicability across ontogeny have been the subject of considerable discussion (Nijhout and German 2012). In particular, Nijhout and colleagues showed that simple power-law relationships may fail to adequately describe allometric patterns during early development, because differences in the timing and kinetics of growth among tissues and organs can produce deviations from a simple scaling relationship. Our results are consistent with this broader observation, although no tissue-specific differences were found in the developmental patterns in the companion paper (Amici et al. 2026).

### The additive constant and early development

What explains the disruption of scale invariance caused by the constant term? One possibility is that early rodent development does not scale proportionally with later milestones when mapped onto human developmental timelines. The temporal dynamics governing early and late developmental stages may not adhere to the same scaling relationship, resulting in a systematic deviation from a simple scale-invariant power law. Closely related species tend to share highly similar developmental programs, see the companion paper. As species diverge evolutionarily, changes accumulate in their developmental milestones timeline. Also these changes do not occur uniformly across ontogeny; rather, they are minimal during early stages and compound as development progresses. It has been more than a century since Haeckel proposed his ideas on the relationship between ontogeny and phylogeny (Levit and Hossfeld 2019). These ideas subsequently contributed to the development of an important field of evolutionary biology, now known as evolutionary developmental biology (evo-devo). Although the relationship between ontogeny and phylogeny is complex, early developmental stages tend to be more conserved across species than later stages. In other words, organisms tend to resemble one another more closely during the early stages of development and progressively diverge as development proceeds. All organisms begin development as a single cell, and the developmental processes immediately following fertilization are relatively similar, at least among deuterostomes. As development proceeds, however, differences in growth and body size, together with their associated geometric and physiological constraints, progressively accumulate and become increasingly evident (Piasecka et al. 2013; Vázquez-Borsetti 2024). It is therefore reasonable to hypothesize that the influence of scaling relationships becomes more pronounced as development progresses. Such differences in the timing of developmental processes across species are encompassed by the concept of heterochrony, which refers to evolutionary changes in the timing or rate of developmental processes (Hall 2012). Also, a contrasting phenomenon may begin to influence development following fertilization, as these trajectories become increasingly constrained by geometric and physiological factors that, at least among mammals, cannot be easily escaped (West et al. 2002). The translation of ontogenetic time across species cannot assume a constant body-mass relationship from fertilization to adulthood. Throughout development, this relationship is inherently variable and diverges progressively, limiting the direct application of quarter-power scaling laws (West et al. 2001). This divergence in developmental trajectories may not be random, but rather amplified along at least three biological axes: the magnitude of allometric differences, phylogenetic distance, and adaptive strategies (i.e., reproductive investment). The integration of these dynamic factors may therefore provide a framework for explaining why the heterochrony observed empirically differs from traditional static allometry. In this context, incorporating an offset may be necessary because a pure power-law relationship would predict the same scaling differences between species throughout development, whereas developmental trajectories may initially be more similar and progressively diverge as differences in body size, mass, and their associated constraints accumulate. From this perspective, the inclusion of an offset may account for an early developmental phase in which species trajectories remain relatively similar, while allowing the scaling relationship to progressively diverge as development proceeds. In this sense, the offset can be interpreted as a correction for the departure from a pure scale-invariant relationship during early development, rather than as evidence for distinct scaling regimes. Another hypothesis is that the constant reflects a feature of comparative developmental biology that has thus far remained concealed. Although the present study does not provide a definitive answer, this remains a compelling avenue for future research. The results of the present study suggest that the relationship between early and late development exhibits a discernible structure that is not adequately described by a linear transformation, but rather follows a quarter-power law with an additive offset. This finding raises the possibility that the temporal relationship between different ontogenetic stages is part of a broader biological scaling pattern. In summary, the selection of the 1/4 exponent model with an additive constant as the definitive model was based on three main considerations. First, from a theoretical perspective, the 1/4 exponent is consistent with proposed scaling principles, providing the model with a biological rationale rather than treating it solely as an empirical curve-fitting approach. Second, the principle of parsimony favors this structure because it requires one fewer parameter than models in which the exponent is freely estimated. Third, constraining the exponent to 1/4 provided a good fit in the individual species analyses and improved the stability of the joint model when both species were analyzed using dummy variables, facilitating the estimation of interspecific effects. Notably, both in the present study and in our companion paper, this model structure provided a useful transformation of developmental time and substantially reduced the nonlinearity observed in the original data. Overall, the quarter-power model with an additive constant provides a parsimonious and theoretically motivated framework for describing developmental timing across humans, mice, and rats.

## AI use statement

The dataset was compiled and curated entirely manually, without the use of artificial intelligence tools for data extraction or curation. AI-assisted tools were used to support code development and to improve the clarity and language of the manuscript. All AI-assisted outputs were reviewed and validated by the author, who take full responsibility for the content of the dataset, the analyses, and the manuscript.

## Data Availability

The code used for data processing, statistical analyses, and figure generation, is openly available in the GitHub repository https://github.com/Vazquez-Borsetti/quarter-power-C-human-rodent-translation. The dataset is permanently archived with a citable DOI in Zenodo: https://doi.org/10.5281/zenodo.23244347. The repository contain the data used in the analyses and the code required to reproduce the reported results.

